# Response to a Synthetic Pheromone Source by OX4319L, a Self-Limiting Diamondback Moth (Lepidoptera: Plutellidae) Strain, and Field Dispersal Characteristics of Its Progenitor Strain

**DOI:** 10.1101/501924

**Authors:** Michael Bolton, Hilda L. Collins, Tracey Chapman, Neil I. Morrison, Stefan J. Long, Charles E. Linn, Anthony M. Shelton

**Affiliations:** School of Biological Sciences, University of East Anglia, Norwich Research Park, Norwich, NR4 7TJ, UK; Department of Entomology, Cornell University, New York State Agricultural Experiment Station, Cornell AgriTech, Barton Laboratory, 630 West North Street, Geneva, New York 14456, USA; Oxitec Ltd, 71 Innovation Drive, Milton Park, Abingdon, Oxfordshire, OX14 4RX, UK

**Keywords:** Pheromone response, field dispersal, trapping, transgenic insects, wind tunnel

## Abstract

The Diamondback Moth, *Plutella xylostella* (L.) (Lepidoptera: Plutellidae), is a global pest that infests vegetable and field crops within the *Brassica* family. A genetically engineered strain of *P. xylostella*, OX4319L, carrying a ‘self-limiting’ gene, has shown potential for managing *P. xylostella* populations, using sustained releases of OX4319L male moths. In order for such a strain to provide control, the transgenic individuals must exhibit attraction to female *P. xylostella* sex pheromone and adequate dispersal in the field. In this study, we tested these key traits. First, we compared the responses of the OX4319L male moths to a synthetic female sex pheromone source in wind tunnel trials to those of males from three other strains. We found that OX4319L males responded comparably to strains of non-engineered males, with all males flying upwind towards the pheromone source. Second, we used mark-release-recapture studies of a wildtype *P. xylostella* strain, from which the OX4319L strain was originally developed, to assess dispersal under field conditions. Released males were recaptured using both pheromone-baited and passive traps within a 2.83 ha circular cabbage field, with a recapture rate of 7.93%. Males were recaptured up to the boundary of the field at 95 m from the central release point. The median dispersal of males was 14 m. These results showed the progenitor strain of OX4319L retained its ability to disperse within a host field. The results of these experiments are discussed in relation to the potential for the effective use of engineered male-selecting *P. xylostella* strains under field conditions.

## INTRODUCTION

The diamondback moth, *Plutella xylostella* (L.) (Lepidoptera: Plutellidae), is a global pest of plants within the *Brassica* family that includes vegetables (e.g. broccoli, cauliflower, mustard, and cabbage) and field crops (e.g. canola). *P. xylostella* -related crop damage and pest management are estimated to cost US $1-5 billion dollars annually (Talekar and Shelton 1993, Zalucki et al. 2012). Although a considerable amount of work has been devoted to various management strategies (Chisholm et al. 1983, Talekar and Shelton 1993, Metz et al 1995, Shelton and Nault 2004, Musser et al. 2005, Badenes-Perez et al. 2005, Hoy et al. 2007, Zalucki et al. 2012, Philips et al., 2014), much of the control effort currently relies primarily on the use of synthetic and biological insecticides. This has led to resistance to almost every class of insecticide, including the microbial insecticide *Bacillus thuringiensis*, and relatively new insecticides such as the anthranilic diamide, chlorantraniliprole (Tabashnik et al. 1990, Talekar and Shelton 1993, Zhao et al. 2006, Wang and Wu 2012). The ability to control *P. xylostella* is becoming increasingly difficult and novel approaches are needed.

The self-limiting pest management approach involves the release of male insects genetically engineered with a gene that prevents survival of female offspring to the reproductive stage. The self-limiting *P. xylostella* strain, OX4319L, was developed by Oxitec Ltd and carries a male-selecting gene that utilizes sequences from the sex determination gene *doublesex* (*dsx*). The gene expresses sex-alternate splicing, to engineer female-specific expression of the self-limiting gene (Jin et al. 2012). This system prevents survival of female offspring beyond the larval stage and allows for production of male-only cohorts of self-limiting moths. After their release, males mate with pest females, leading to a reduction in the number of female offspring in the next generation, thereby suppressing *P. xylostella* populations. To facilitate the rearing of large numbers of males for release within DBM production facilities, the expression of female-specific *dsx* within the OX4319L strain is repressed by the addition of tetracycline, or suitable analogues, into the larval feed. OX4319L also expresses the fluorescent protein, DsRed, to permit the effective monitoring of the presence of this strain in the field. Such genetic systems have been successfully developed in a number of insect species such as in *P. xylostella* (Jin et al. 2013), medfly (*Ceratitis capitata*) (Fu et al. 2007), and olive fly (*Bactrocera oleae*) (Ant et al. 2012).

The OX4319L *P. xylostella* strain has undergone extensive laboratory and glasshouse testing to assess fitness-related performance traits such as mating competitiveness, its ability to suppress *P. xylostella* populations, and to identify any fitness costs related to the transgene insertion itself (Harvey-Samuel et al. 2014). These experiments have shown that OX4319L can cause population suppression of wildtype individuals and it also has the potential to reduce the development of insecticide resistance (Harvey-Samuel et al. 2015). However, the performance of this strain in two other key respects, pheromone attraction and field dispersal, has not yet been tested.

These traits are important to assess because future releases of self-limiting *P. xylostella* strains are likely to use trapping to monitor field releases of moths and to assess the suppression performance of the strain in the field. This is most likely to be achieved using pheromone-baited traps. In addition, the ability of self-limiting males to seek out and mate with wild females will be largely influenced by their ability to respond to its sex pheromone.

The role of sex pheromone communication in lepidopteran species is highly complex and species-specific (Allison and Cardé 2016). Pheromone communication is known to play a key role in mating success and to have an important influence on reproductive isolation and, potentially, speciation. In laboratory experiments, changes in sex pheromone communication represent the basis for divergence among closely related species (Tabata and Ishikawa 2005). Assortative mating within strains of the same insect species with different pheromone components, or different ratios of those pheromone components, has also been described (Zhu et al. 1997, Pélozuelo et al. 2004, Pélozuelo et al. 2007). Environmental factors can have a large influence on changes in pheromone-based communication systems (Svensson et al. 2002, Yang and Du 2003, Pélozuelo et al. 2004, Robbins et al. 2008). It is possible, through differing selection pressures or genetic drift, that populations of *P. xylostella* maintained over many generations in laboratory or factory-rearing conditions could develop distinct behaviors or characteristics that would affect their interactions with wild individuals in an open-field setting or their ability to be monitored by pheromone lures. Such possibilities should be examined in order to evaluate the potential for new control technologies.

One aspect of *P. xylostella* ecology that facilitates its status as one of the globe’s most devastating agricultural pests is its capacity for dispersal and migration. Movement of *P. xylostella* can be divided into two broad categories: long-distance migration and short-distance dispersal (Schowalter 2006). Shorter-distance dispersal would be important for the success of a self-limiting control approach to target *P. xylostella* in the field. A critical aspect of any release program is the ability of released males to disperse a sufficient distance and seek out and mate with wild females. For example, the number and location of male releases needed for control would be dependent upon the flight ability of the released population of males. The mean dispersal distance of wildtype *P. xylostella* in previous mark-release-recapture trials has been measured as 21-35 m when capturing males with ‘active’ pheromone-baited traps, and 14-18 m and 13-24 m when using yellow, sticky bucket ‘passive’ traps to capture males and females, respectively (Mo et al. 2001). Furthermore, in open-field situations 95% of males were recaptured within 102 m of their release site when using pheromone-baited traps, and within 54 m using yellow, sticky bucket traps (Mo et al. 2001).

In this study, we first evaluated the responsiveness of *P. xylostella* strains to a synthetic pheromone to investigate this aspect of mate-seeking by OX4319L males and hence the likely success of using pheromone-baited traps for monitoring or trapping during control programs. This was done by testing the response of OX4319L males to a synthetic pheromone source in a wind tunnel in comparison to that of males from two laboratory-reared strains and males from one recent field-collected strain of *P. xylostella*. We then explored the dispersal characteristics of a long term, laboratory-reared strain in a host field using a mark-release-recapture technique. The laboratory-reared strain used, ‘Vero Beach’, is the progenitor strain of the OX4319L *P. xylostella*. The dispersal characteristics of these *P. xylostella* were assessed using both passive and pheromone-baited traps.

## MATERIALS AND METHODS

### Diamondback moth strains

The *P. xylostella* strains utilized for laboratory experiments were Geneva 88, OX4319L, Georgia, and Vero Beach. Only the Vero Beach strain was used for the field research component of the study.

### History and rearing of strains

The Geneva 88 strain was first colonized in the laboratory in 1988 after being field-collected in Geneva, NY. It has been maintained in the NYSAES laboratories since that date. The Vero Beach strain was acquired by Oxitec Ltd from Syngenta AG, Jealott’s Hill, UK in 2008, and was reared at Oxitec Ltd facilities until sent to NYSAES laboratories in May 2014. The Vero Beach strain was used to engineer the OX4319L transgenic line of *P. xylostella*, and therefore served as the best available non-engineered control strain for OX4319L. These three populations were reared using the same non-tetracycline larval diet as the OX4319L population (Shelton et al. 1991).

A strain field-collected from Omega Co., GA in May 2014, hereafter referred to as the ‘Georgia’ strain, was also reared in the NYSAES facility. This population was reared on broccoli plants in large cages within glasshouses with a light cycle of 16 h:8 h light to dark, at 25 °C. Cages were approximately 2 m^3^ and contained 8-10 potted broccoli plants at any one time. The moth population fluctuated across generations and was not routinely monitored. Fresh broccoli plants were introduced to the cages weekly to maintain a constant source of food to support larval development.

Adult moths from the Georgia and Geneva 88 strains were housed in smaller, cylindrical cages (20×30 cm) of approximately 500 individuals and fed with sugar water solution. Eggs were collected using sections of aluminum foil painted with cabbage juice residue as an attractant (Shelton et al. 1991). OX4319L moths were housed in Bugdorm cages (30×30×30 cm; BioQuip, Rancho Dominguez, CA) containing approximately 2000 individuals and supplied with a solution for adult feeding (7.5% (wt/vol) sucrose with 0.01% tetracycline). All laboratory strains were maintained at 25 °C with 50% relative humidity (RH), on a 16 h:8 h light to dark cycle, in controlled environment facilities.

The OX4319L strain has been genetically engineered to incorporate a tetracycline-repressible, male-selecting (female-lethal) genetic construct into the *P. xylostella* genome (Jin et al. 2013). For colony maintenance, provision of chlortetracycline hydrochloride (100 μg/L) in the larval diet allowed both males and females to survive and maintain a stock population. To provide the males used for experiments, eggs were seeded onto larval media containing no chlortetracycline hydrochloride, leading to production of male-only cohorts of adult moths. All larvae were reared in a constant-temperature chamber at 25 °C and 50% RH, with a 16 h:8 h light to dark cycle, so that “dusk” occurred in the afternoon.

### Wind tunnel facility

Observations were made in a wind tunnel using the set-up described by Miller and Roelofs (1978). Briefly, the wind tunnel was approximately 2 m in length and generated an air speed of 0.5 m/s throughout. The tunnel floor was static but supplemented with randomly spaced artificial green paper circles (varying 7.6-12.7 cm in diameter) to provide males with a contrasting, non-directional, visual field to aid their optomotor upwind flight response.

### Response of strains to synthetic pheromone in a wind tunnel

Pupae were collected from all four strains, the silk case removed by hand and the pupae then kept individually in 29.6 mL lidded, plastic cups. Males were housed in the constant-temperature chamber (25 °C and 50% RH) for 24-48 h post-eclosion before any flight trials and supplied with a sugar water. Male moths were moved from their growth chamber 1 h before the dark cycle started into the environmental controlled room housing the wind tunnel and transferred into mesh cages from which flight trials could be conducted. Males were left for 1 h in the room housing the wind tunnel prior to experimentation to allow them to acclimatize to conditions in the facility. After lights were turned off, males were given another 15 m to acclimatize to dark conditions (10-15 lux) before any trials were initiated. During this time, the synthetic pheromone lure (ISCAlure-Xylostella, ISCA Technologies, Riverside, CA) was added to the upwind end of the wind tunnel and placed on a copper tube at the end of a metal stand. Male moths were placed in the downstream end of the wind tunnel individually and allowed to exit freely from their mesh cages. These males were monitored and assessed against three behavioral checkpoints: take flight (initiate flight); fly upwind (oriented flights facing upwind and within the odor plume); close (fly to within 10 cm of the pheromone source).

Multiple males from each strain were used in the flight trials per day, with each male being used only once. The procedure was carried out over several days during a period of 3 wks. The number of males used from each strain varied slightly among days due to time constraints of “dusk”-time testing and logistical constraints of rearing sufficient numbers of males from laboratory and wild-caught strains. The trials were continued until a minimum of 30 males had been tested for each strain.

### Field dispersal of the Vero Beach strain

The field site used for releases was on the NYSAES Research Farm, Geneva, NY. Within a 2.83 ha circular cabbage-planted field, 48 traps (Figure 1) were positioned in eight concentric circles around a central release point (Figure 2). Traps consisted of an inverted 355 ml Styrofoam cup with a 3.3 cm wide plastic rim at the base. The base and cup were coated with Tanglefoot^®^ (Olson Products Inc., Medina, OH) that were secured to fiberglass poles approximately 0.5 m above the ground (just above the plant canopy) with a pheromone lure (if baited) attached approximately 2 cm above the trap (Musser et al. 2005). Forty-eight traps were placed in the field in concentric circles at the following distances from the release site: 2 traps at 7 m, 4 traps at 14 m, 8 traps at 21 m, 10 traps at 28 m, 12 traps at 35 m, 4 traps at 55 m, 4 traps at 75 m and 4 traps at 95 m. Traps in a given concentric circle were equidistant from each other. The purpose of the inner circles was to provide data on the closer movement of released moths. The purpose of the more distant rings was to provide data on the upper limit of moth dispersal within the circular field.

**Figure 1.**
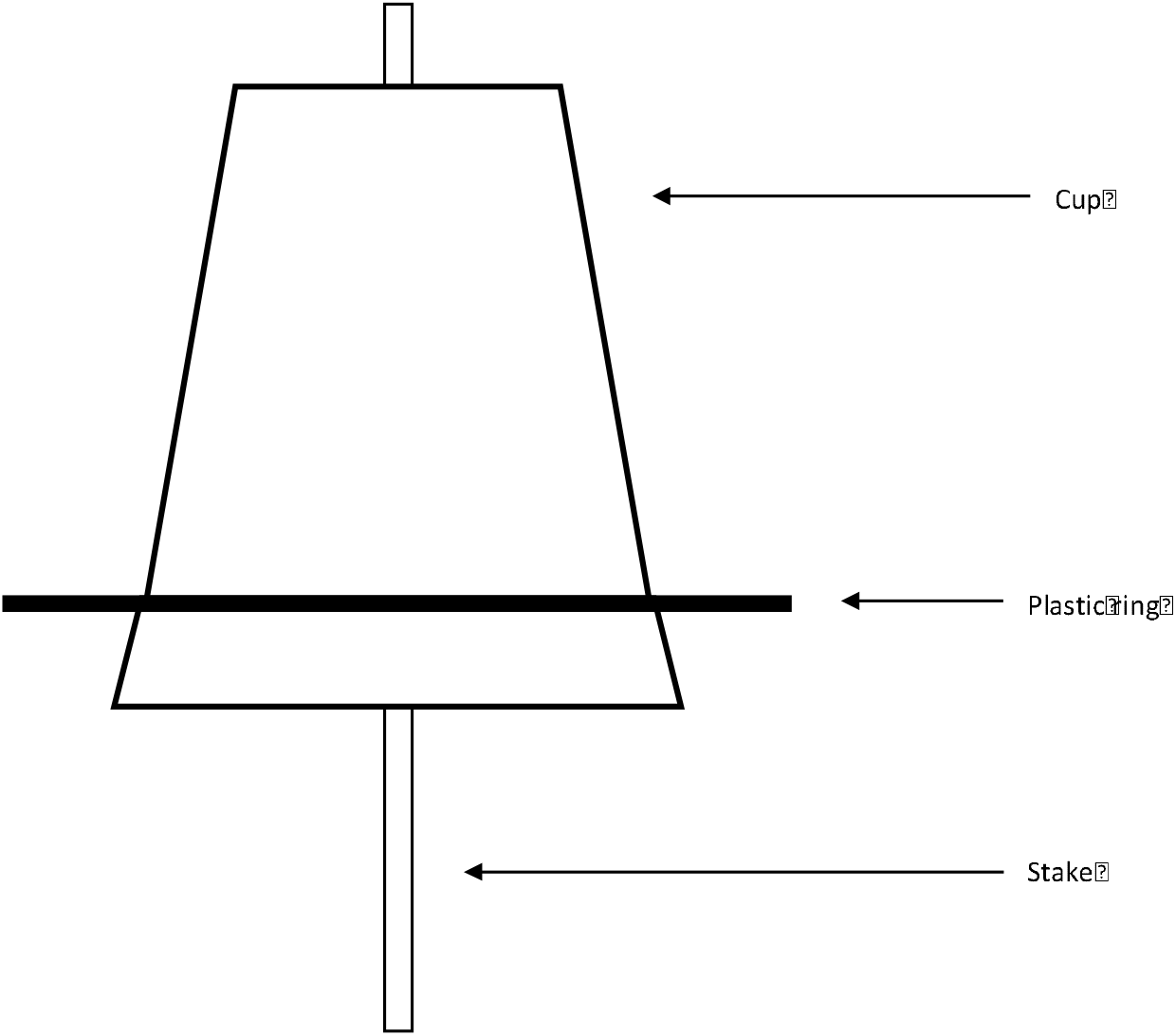
Trap design comprising an upturned cup with plastic ring at the base, all of which was coated with Tanglefoot, based on the design of Musser et al. (2005). Where appropriate, pheromone lures were attached at the top of the cup, secured to the stake with a bulldog clip.

**Figure 2.**
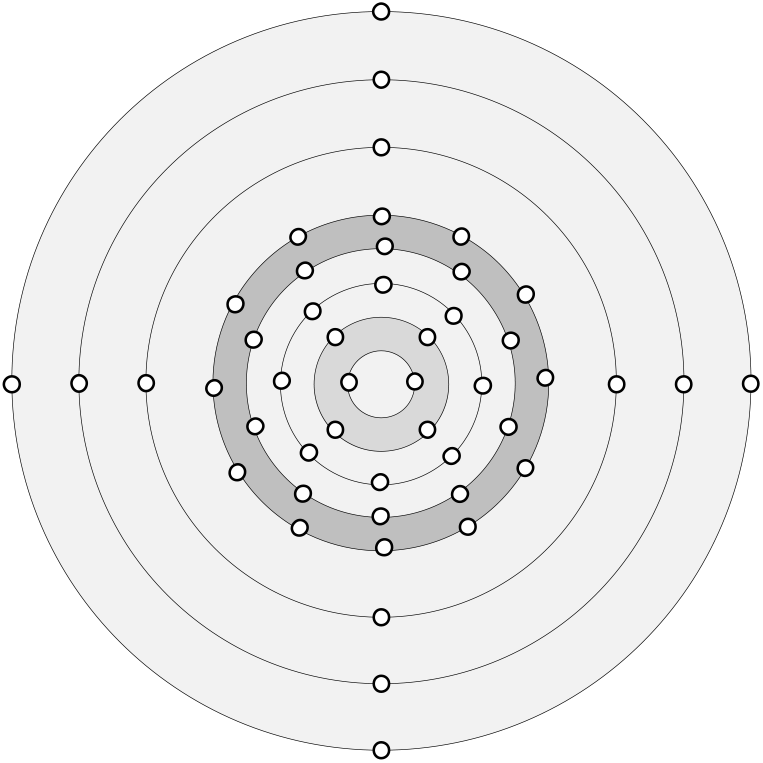
Schematic diagram of trap layout in the circular cabbage-planted field. White circles illustrate trap positions in concentric rings around a central release point. The inner five rings, up to and including the grey ring, were spaced 7 m apart, encompassing a total radius of 35 m from the central release point (7 m, 14 m, 21 m, 28 m, and 35 m from the release point respectively). The outer three rings were spaced 20 m apart (55 m, 75 m, and 95 m from the release point respectively).

Half of the sticky traps placed in the field were also baited with a pheromone lure to assess their attractiveness to Vero Beach moths, the progenitor of the OX4319L moths. At each trap distance, half of the traps were baited with a pheromone lure in an alternating fashion. For the traps at 7 m, 14 m and 21 m, the first trap at each distance was assigned the same treatment (baited or not baited with pheromone) because the first trap at 14 m was offset from the line formed between the 1^st^ traps at 7 m and 21 m by an angle of 45 degrees. The first trap at 28 m was assigned the opposite treatment of the first trap at 21 m. The first trap at the next distance was assigned the opposite configuration (not baited if the previous was baited) for distances greater than 21 m, i.e. if trap 1 in the 7 m ring was baited with a pheromone lure, then in the 28 m ring trap 1 was not baited, trap 2 was baited and so on. For the releases monitored during August, trap 1 at 7 m was baited with pheromone. For the releases monitored during September, trap 1 at 7 m was not baited with pheromone. This adjustment was to negate impacts of prevailing wind direction.

The Vero Beach populations of mixed sex were maintained in cylindrical cages (20 × 30 cm) containing approximately 500 adult moths. Eggs were collected from these populations on aluminum foil painted with cabbage juice as an attractant (Shelton et al. 1991). These eggs were seeded onto non-tetracycline larval diet and allowed to develop until pupation. Pupae were then transferred into a Bugdorm cage (30×30×30 cm) and allowed to eclose.

Upon eclosion, male moths were aspirated out of the Bugdorm containers every 3 h over a period of 48 h and moved to cylindrical release cages, each containing up to 250 male moths and a sugar water source. All females were discarded. The collected male cohorts were grouped according to the 24 h window in which they were collected, resulting in a pair of releases on consecutive days. Males were dusted with a fluorescent powder (Day-Glo Corp., Cleveland, OH), with a different color used for each day of release to facilitate recognition of the release day in the recaptured individuals. Once collected and dusted, males were left for between 12 h and 24 h before release to ensure all males were at least 24 h old.

Release cages were transported to the center of the cabbage field 1 h before dusk and left to acclimatize after a 10 min journey by car. At dusk, the lids and bases were removed from the cylindrical release cages and moths were allowed to freely disperse into the cabbage field. Cages were left at the release point until all moths had exited. This release procedure was repeated the next day with the second cohort of collected males.

Traps were checked the morning after any releases took place and every 2 d thereafter. Any traps containing moths were removed, taken back to the laboratory for moth identification via scoring for the fluorescent dust under UV light, and replaced with a fresh trap. The number of males of each dusted color was scored for each collected trap. The presence and color of fluorescent powder indicated whether it was a released male and, if so, the release day. Trapping continued until no dusted moths were recaptured. This release experiment protocol was monitored during two time periods: one during August and the other during September 2014. For the first release period, 638 males were released on July 31 followed by a release of 355 males on August 1. For the second release period, 500 males were released on September 7 and another 500 males were released on September 8. The mean distance travelled (MDT) for moths recaptured on sticky traps with and without pheromone lures was calculated according to the method of Morris et al. (1991) for each release.

### Data Analysis

Data analysis was performed using R (R Core Team 2013). Analysis of behavior in the wind tunnel experiments was performed using Fisher’s exact tests.

The mean distance travelled was analyzed using a mixed model with the square root of the daily MDT as the response variable and trap type (with or without pheromone) and days post-release and their interaction (full factorial) as fixed effects and release number as a random effect. Means were separated using Student’s t-test at alpha=0.05.

## RESULTS

### Response of strains to synthetic pheromone in a wind tunnel

Wind tunnel behaviors were analyzed independently by assessing the proportion of males completing each behavior in the different *P. xylostella* strains (Figure 3). No significant differences were observed between strains in any of the behaviors: Take flight (p = 0.3298), Fly upwind (p = 0.4647), and Close (p = 0.6117). These results suggest that the strains did not show significantly different responses, relative to each other, to the synthetic pheromone lure placed in the wind tunnel experiment.

**Figure 3.**
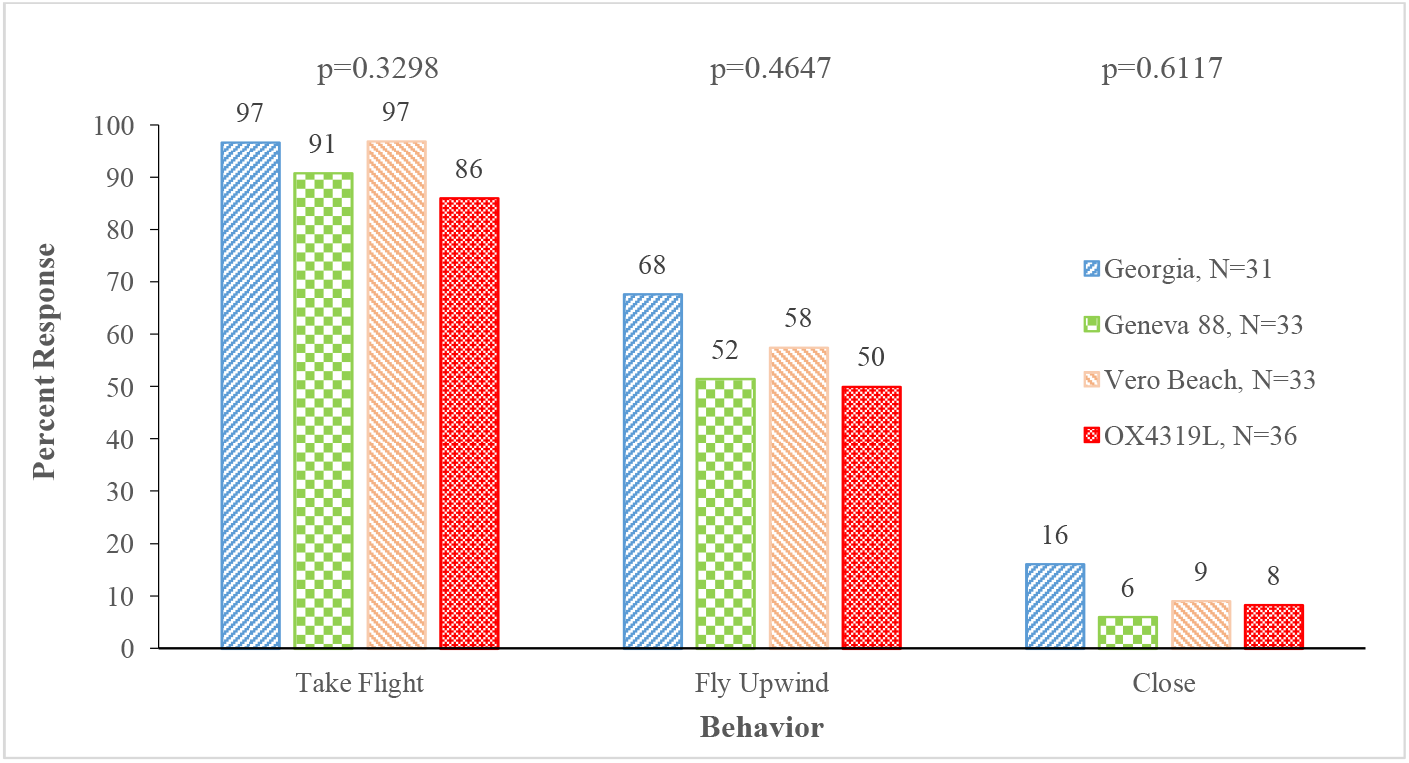
Behavioral responses of male moths from different *P. xylostella* strains to a synthetic pheromone lure in a wind tunnel. The data shown are the percentage of each strain completing each of the behavioral checkpoints listed on the x-axis (OX4319L, n = 36; Vero Beach, n = 33; Geneva 88, n = 33; Georgia, n = 31). These checkpoints were: *Take Flight*, whether a male took flight during the flight trial; *Upwind*, whether a male moved upwind towards the synthetic pheromone source; and *Close*, whether a male reached within 10 cm of the pheromone lure. p-values for each behavior were determined using Fisher’s exact test with α=0.05.

### Field dispersal of the Vero Beach strain

Of the total number of Vero Beach male moths released (n = 1993), a recapture rate of 7.93% was recorded across all releases (Table 1). Traps without pheromone did not detect moth dispersal beyond 28 m, whereas traps with pheromone detected moths at every distance out to 95 m.

**Table 1.**
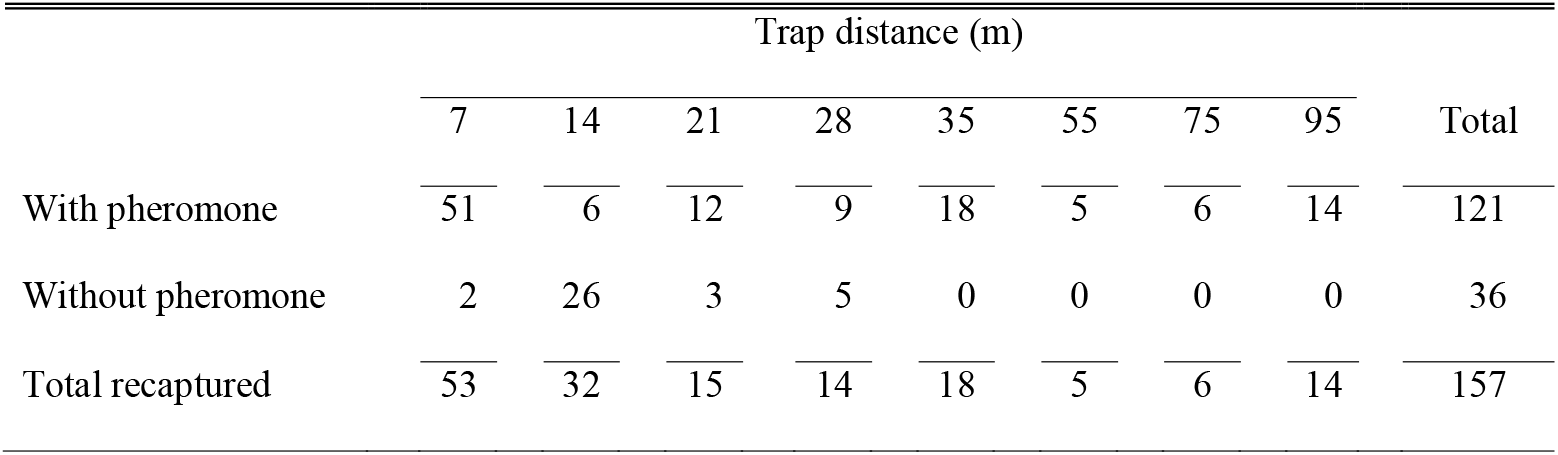
Number of *P. xylostella* recaptured on sticky traps with and without pheromone lures.

|  | Trap distance (m) |  |  |  |  |  |  |  | Total |
| --- | --- | --- | --- | --- | --- | --- | --- | --- | --- |
|  | 7 | 14 | 21 | 28 | 35 | 55 | 75 | 95 |  |
| With pheromone | 51 | 6 | 12 | 9 | 18 | 5 | 6 | 14 | 121 |
| Without pheromone | 2 | 26 | 3 | 5 | 0 | 0 | 0 | 0 | 36 |
| Total recaptured | 53 | 32 | 15 | 14 | 18 | 5 | 6 | 14 | 157 |

Both types of traps were able to detect moths up to 7 d post release (Table 2). Traps with pheromone lures detected significantly greater dispersal than traps without pheromone lures each day throughout the release period. The overall mean distance travelled by moths recaptured on traps with pheromone lures was 54.6 m compared to 17.8 m on traps without pheromone lures (Table 2). Also, the mean distance travelled increased over the 7-d monitoring period indicating that moths dispersed continually throughout this period. No trapping was carried out outside of the circular cabbage-planted field, so the maximum dispersal of the released moths outside of the host range tested is unknown.

**Table 2.**
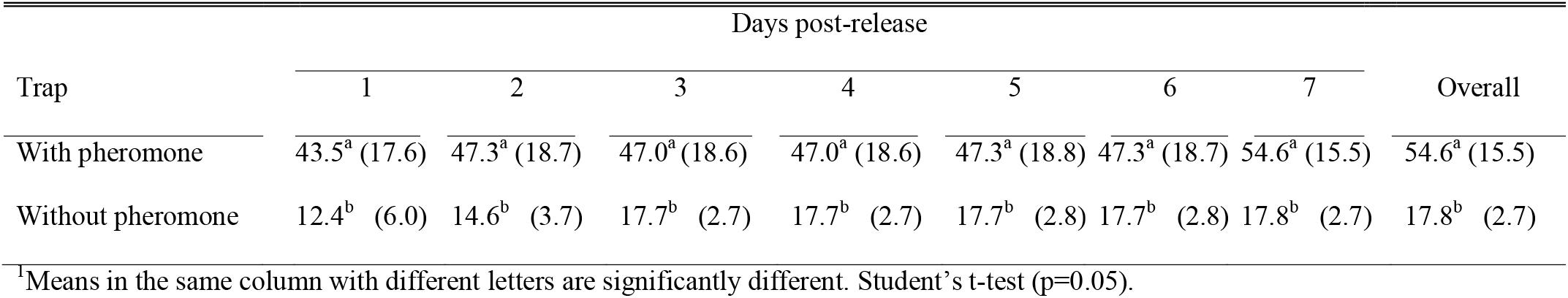
Mean^1^ (SE) distance travelled for *P. xylostella* recaptured on sticky traps with and without pheromone lures.

| Trap | Days post-release |  |  |  |  |  |  | Overall |
| --- | --- | --- | --- | --- | --- | --- | --- | --- |
|  | 1 | 2 | 3 | 4 | 5 | 6 | 7 |  |
| With pheromone | 43.5<sup>a</sup> (17.6) | 47.3<sup>a</sup> (18.7) | 47.0<sup>a</sup> (18.6) | 47.0<sup>a</sup> (18.6) | 47.3<sup>a</sup> (18.8) | 47.3<sup>a</sup> (18.7) | 54.6<sup>a</sup> (15.5) | 54.6<sup>a</sup> (15.5) |
| Without pheromone | 12.4<sup>b</sup> (6.0) | 14.6<sup>b</sup> (3.7) | 17.7<sup>b</sup> (2.7) | 17.7<sup>b</sup> (2.7) | 17.7<sup>b</sup> (2.8) | 17.7<sup>b</sup> (2.8) | 17.8<sup>b</sup> (2.7) | 17.8<sup>b</sup> (2.7) |

<sup>1</sup>Means in the same column with different letters are significantly different. Student's t-test (p=0.05).

Of those males recaptured, 75% (n = 119) were recaptured using traps baited with synthetic pheromone lures. Passive traps were only effective up to a distance of 28 m, after which recapture of moths was observed only on the pheromone-baited traps. A number of non-dusted moths (n = 363) were captured on both passive and pheromone-baited traps across the two releases, indicating the presence of a background population of wild *P. xylostella* present within the cabbage field.

## DISCUSSION

In the wind tunnel, males were assessed against three critical behavioral checkpoints and no significant differences among strains at any of the checkpoints tested were found. Most importantly, the key checkpoints of ‘take flight’ and ‘fly upwind’ showed no significant differences between the OX4319L and other strains. This suggests that, in future field applications, OX4319L males will show comparable responses to the female sex pheromone as wild-type counterparts. The last behavioral checkpoint (‘close’) had the lowest proportion and no males landed on or made contact with the pheromone lure. This was probably due to the very high pheromone concentrations given off by the lure. The lures used in this experiment were commercial lures, which are designed to give off high levels of pheromones to attract males from a wide area. The higher relative concentrations that these commercial lures would have produced inside the wind tunnel could cause males to arrest their upwind oriented flight approach to the pheromone source (Linn and Roelofs. 1983).

While this experiment suggests that the ability of OX4319L males to react to pheromone sources was not diminished in comparison to other strains, it does not answer the question of whether OX4319L males can achieve mating success or paternity once they have actively responded to a true pheromone source emitted by wild females.

During the field dispersal experiments, males from a long-term, laboratory-reared strain (Vero Beach), which was the progenitor of the OX4319L, were capable of flight and dispersal within a field of cabbage. Males were recaptured at all distances where traps were placed within the circular field up to the maximum distance of 95 m. Of the moths recaptured, the mean and median recapture distances for Vero Beach males were 26.8 m and 14 m, respectively. These values are consistent with the findings of Mo et al. (2001), who reported the mean dispersal distance of male *P. xylostella* to be 21-35 m and 14-18 m when using pheromone-baited and passive traps, respectively. The data also agree with the findings of Musser et. al (2005) that *P. xylostella* adults released from a central point are caught in higher numbers near the release site but do tend to disperse over time.

Males were only recaptured at distances greater than 28 m using traps baited with a synthetic pheromone lure. This is consistent with findings from the literature, and the finding that pheromone traps can actively draw in males from outside the release site (Mo et al. 2001, Mo et al. 2003). This suggests that the presence of pheromone-baited traps may affect the behavior of male moths and therefore may not be the best method of studying steady-state dispersal characteristics. The use of passive traps might provide a better representation of the natural dispersal characteristics of males of any given strain. However, when trapping methods are being used to monitor *P. xylostella* pest numbers, it may be beneficial to recapture as many individuals as possible using a pheromone-baited trap. Overall, the experiments described here demonstrate that laboratory-reared strains can be readily tracked after release using mark-release-recapture techniques, using fluorescent dusts with passive and active trapping methods. Moreover, the data provide valuable information about the ability of the OX4319L to respond to a synthetic pheromone source. This suggests they may also be attracted to pest females in the field and that pheromones may be used as an easy tracking method for OX4319L in the field.

Most importantly, these data also show the ability of the progenitor strain of OX4319L to disperse when released in male-only cohorts, as would be required for application of OX4319L against *P. xylostella*. Data on the average distance dispersed is an important factor when planning such releases, as adequate coverage of self-limiting males is key to providing a sufficient suppressive effect on the pest population. For mating-based pest management, long distance dispersal is advantageous as it allows for fewer release points, providing more economical deployment of the self-limiting males. However, in scenarios of localized outbreaks, a more concentrated release might be a better option.

The experiments described here further demonstrate the promise of this self-limiting technology for pest management of *P. xylostella*. These data suggest that the OX4319L strain of *P. xylostella* is able to detect and respond to a synthetic pheromone source, comparable to its wildtype counterparts. The behavior of this strain in a wind tunnel, combined with similar dispersal behavior of the progenitor strain of OX4319L within a host field, are promising signs that OX4319L would be able to disperse and find females in the field. Further research should be directed at their ability to mate with females in the field and suppress an emerging population, as has been demonstrated in greenhouses (Harvey-Samuel et al. 2015).

## ACKNOWLEDGMENTS

We thank the UK National Environment Research Council CASE studentships for doctoral funding for M. Bolton. Funding for this study was provided by Oxitec/Intrexon as a sponsored grant that precluded their influence on the study and its outcome.

